# Whole genome sequences of 23 species from the *Drosophila montium* species group (Diptera: Drosophilidae): A resource for testing evolutionary hypotheses

**DOI:** 10.1101/861005

**Authors:** Michael J. Bronski, Ciera C. Martinez, Holli A. Weld, Michael B. Eisen

## Abstract

Large groups of species with well-defined phylogenies are excellent systems for testing evolutionary hypotheses. In this paper, we describe the creation of a comparative genomic resource consisting of 23 genomes from the species-rich *Drosophila montium* species group, 22 of which are presented here for the first time. The *montium* group is uniquely positioned for comparative studies. Within the *montium* clade, evolutionary distances are such that large numbers of sequences can be accurately aligned while also recovering strong signals of divergence; and the distance between the *montium* group and *D. melanogaster* is short enough so that orthologous sequence can be readily identified. All genomes were assembled from a single, small-insert library using MaSuRCA, before going through an extensive post-assembly pipeline. Estimated genome sizes within the *montium* group range from 155 Mb to 223 Mb (mean=196 Mb). The absence of long-distance information during the assembly process resulted in fragmented assemblies, with the scaffold NG50s varying widely based on repeat content and sample heterozygosity (min=18 kb, max=390 kb, mean=74 kb). The total scaffold length for most assemblies is also shorter than the estimated genome size, typically by 5 - 15 %. However, subsequent analysis showed that our assemblies are highly complete. Despite large differences in contiguity, all assemblies contain at least 96 % of known single-copy Dipteran genes (BUSCOs, n=2,799). Similarly, by aligning our assemblies to the *D. melanogaster* genome and remapping coordinates for a large set of transcriptional enhancers (n=3,457), we showed that each *montium* assembly contains orthologs for at least 91 % of *D. melanogaster* enhancers. Importantly, the genic and enhancer contents of our assemblies are comparable to that of far more contiguous *Drosophila* assemblies. The alignment of our own *D. serrata* assembly to a previously published PacBio *D. serrata* assembly also showed that our longest scaffolds (up to 1 Mb) are free of large-scale misassemblies. Our genome assemblies are a valuable resource that can be used to further resolve the *montium* group phylogeny; study the evolution of protein-coding genes and *cis*-regulatory sequences; and determine the genetic basis of ecological and behavioral adaptations.

## Introduction

Large groups of closely related species with well-defined phylogenetic relationships are invaluable resources with which to investigate evolutionary processes [1]. Previous comparative genomic studies in *Drosophila* have included twelve species spanning the entire *Drosophila* lineage [1]. Taking into account the short generation time of *Drosophila*, the evolutionary divergence of this sample size space exceeds that of the entire mammalian radiation [1,2]. Subsequent sequencing efforts added eight genomes at intermediate evolutionary distances from *D. melanogaster* [3]. *While these data sets have provided extraordinary insight into Drosophila* evolution, they also pose unique challenges. As phylogenetic distance from *D. melanogaster* increases, it becomes more difficult to identify orthologous sequence [3]; and multi-species alignments with divergent sequences can be sensitive to alignment error, especially for small features such as transcription factor binding sites [2,4]. Accordingly, a data set is needed where 1) distances from *D. melanogaster* are short enough so that orthologous sequence can be readily identified, and 2) species are closely related enough such that sequence similarity produces accurate alignments, but distantly related enough to recover a strong signal of sequence divergence. In this paper, we describe the creation of such a resource by assembling 23 genomes from the *Drosophila montium* species group. Genomes for 22 of these species are presented here for the first time.

The *Drosophila montium* species group [5–7] *is the largest group in the subgenus Sophophora*. This species-rich clade contains 94 species [8,9] *currently divided into seven subgroups [7]. The montium* group diverged from *D. melanogaster* roughly 28 million years ago (mya) [10], and the most recent common ancestor of all *montium* species lived approximately 19 mya [7]. Members of the *montium* group are distributed across Africa, South Asia, South-East Asia, East Asia, and Oceania [7]. More than 40 species are currently available in culture, and the list continues to grow. Species from the *montium* group have been used to study a variety of evolutionary, ecological, and behavioral questions, including the genetic basis of female-limited color polymorphism [11], cold and desiccation resistance [12], adaptation to drought stress [13], and courtship behavior [14,15]. Previous phylogenetic reconstructions of the *montium* group - typically based on small numbers of genes - have produced incongruent trees, although recent reconstructions are generally well-resolved and congruent [5,11,14–18].

Two *montium* genomes have already been assembled. The *D. kikkawai* genome [3] was sequenced to a depth of 182x coverage using a combination of 454 and Illumina technology. This produced a 164 Mb assembly with a scaffold N50 of 904 kb. The *D. serrata* genome [19] was sequenced to a depth of 63x coverage using PacBio long-reads. It yielded a 198 Mb assembly with a contig N50 of 943 kb. While these approaches generated high-quality draft assemblies, the associated costs preclude sequencing dozens of *montium* species this way.

Our goal therefore was to assemble dozens of *montium* genomes in a cost-effective way, while also producing assemblies of sufficient quality and completeness to study protein-coding genes and non-coding sequences genome-wide. In this paper, we describe the sequencing and assembly of 23 *montium* genomes. While our assemblies are relatively fragmented, our analysis shows they are also highly complete. All assemblies contain high percentages of known genes and enhancers, and by these measures, they are indistinguishable from far more contiguous *Drosophila* assemblies. Going forward, our assemblies will be a valuable resource that can be used to further resolve the *montium* group phylogeny; study the evolution of protein-coding genes and *cis*-regulatory sequences; and determine the genetic basis of ecological and behavioral adaptations.

## Results and Discussion

### Genome size estimates and assembly statistics

To assemble dozens of genomes in a cost-effective way, we sequenced a single, small-insert (350 bp), PCR-free, library to roughly 35x coverage for each species. The genomes were assembled using the Maryland Super Read Cabog Assembler (MaSuRCA), which combines de Bruijn graph and overlap-layout-consensus (OLC) approaches into a novel algorithm based on “super-reads” [20]. The genomes then went through an extensive post-assembly pipeline to further improve the primary assemblies. See the Materials and Methods for an in-depth description of the entire pipeline.

Table 1 reports genome size estimates and assembly statistics (total scaffold length, scaffold / contig NG50, length of longest scaffold / contig, and total gap length) for 23 *montium* species. Genome size estimates are based on the *k*-mer frequency spectrum of the unassembled reads, as calculated by String Graph Assembler (SGA) Preqc [21]. The scaffold / contig NG50 [22,23] is analogous to the well-known N50, but substitutes the estimated genome size for the total assembly length. For example, a scaffold NG50 of 100,000 bp means that 50 % of the estimated genome size is present in scaffolds that are at least 100,000 bp. When this calculation is repeated for all integers from 1 to 100, the result is an “NG graph” [23]. Figure S1 contains NG graphs showing the distribution of scaffold lengths for each *montium* assembly. Table 2 contains additional sample information, including the strain name, coverage, and GC %. The table also reports the frequency of variant and repeat branches in de Bruijn graphs constructed by SGA Preqc [21], which are estimates of heterozygosity and repeat content, respectively.

**Table 1.** Genome size estimates and assembly statistics. Genome size estimates were calculated by SGA Preqc [21] based on the *k*-mer frequency spectrum of the unassembled reads. To calculate the scaffold NG50 [22,23], scaffold lengths were ordered from longest to shortest and then summed, starting with the longest scaffold. The NG50 was the scaffold length that brought the sum above 50 % of the estimated genome size. Contig lengths were estimated by splitting scaffolds on every N (including single Ns). Species are listed in decreasing order of scaffold NG50.

| **Species** | **Est. Genome Size (bp)** | **Total Scaffold Length (bp)** | **Scaffold NG50 (bp)** | **Longest Scaffold (bp)** | **Contig NG50 (bp)** | **Longest Contig (bp)** | **Total Gap Length (bp)** |
| --- | --- | --- | --- | --- | --- | --- | --- |
| *D. kanapiae* | 155,490,160 | 152,203,088 | 389,587 | 2,274,126 | 301,459 | 2,274,126 | 205,953 |
| *D. birchii* | 169,148,727 | 156,593,892 | 211,718 | 1,501,252 | 164,580 | 1,183,551 | 81,207 |
| *D. truncata* | 190,688,284 | 167,897,087 | 95,737 | 830,117 | 73,889 | 827,712 | 375,696 |
| *D. bocki* | 155,095,574 | 151,202,254 | 78,068 | 785,450 | 65,314 | 785,450 | 69,231 |
| *D. bunnanda* | 181,250,127 | 151,823,105 | 76,713 | 1,142,480 | 65,403 | 1,127,760 | 152,423 |
| *D. punjabiensis* | 197,448,094 | 192,339,030 | 72,420 | 1,226,934 | 64,043 | 1,083,757 | 153,372 |
| *D. jambulina* | 179,468,675 | 163,991,206 | 71,637 | 873,064 | 61,348 | 756,687 | 61,365 |
| *D. vulcana* | 209,187,412 | 187,578,810 | 65,096 | 530,507 | 51,774 | 472,464 | 116,296 |
| *D. seguyi* | 206,814,592 | 178,856,532 | 63,109 | 891,413 | 54,123 | 891,413 | 233,653 |
| *D. mayri* | 223,398,425 | 167,807,061 | 62,249 | 2,219,437 | 43,922 | 1,355,909 | 364,662 |
| *D. asahinai* | 216,977,949 | 189,050,820 | 59,266 | 1,052,132 | 50,528 | 904,342 | 75,577 |
| *D. serrata* | 184,673,878 | 159,679,625 | 54,224 | 1,091,401 | 43,626 | 718,797 | 79,019 |
| *D. lacteicornis* | 203,475,870 | 182,681,050 | 53,799 | 1,044,495 | 44,105 | 766,914 | 60,537 |
| *D. pectinifera* | 220,219,034 | 149,209,000 | 52,632 | 528,734 | 41,478 | 467,725 | 58,142 |
| *D. tani* | 194,820,185 | 180,972,673 | 48,517 | 921,527 | 44,341 | 780,901 | 135,927 |
| *D. rufa* | 210,769,271 | 186,167,886 | 43,287 | 498,065 | 38,201 | 498,065 | 80,447 |
| *D. watanabei* | 182,199,997 | 196,825,890 | 40,952 | 1,045,963 | 36,818 | 656,929 | 135,921 |
| *D. auraria* | 220,036,088 | 197,420,731 | 38,365 | 491,046 | 35,679 | 491,046 | 130,216 |
| *D. bakoue* | 219,308,053 | 187,248,584 | 28,924 | 1,045,797 | 27,520 | 598,596 | 58,713 |
| *D. leontia* | 162,918,854 | 164,601,511 | 26,461 | 331,217 | 23,897 | 301,031 | 91,716 |
| *D. burlai* | 198,129,694 | 175,666,184 | 24,417 | 628,960 | 23,536 | 628,960 | 153,793 |
| *D. nikananu* | 217,706,973 | 190,505,469 | 23,001 | 626,542 | 21,180 | 574,375 | 226,904 |
| *D. triauraria* | 217,036,792 | 197,369,186 | 17,513 | 590,840 | 16,493 | 576,941 | 156,570 |

**Table 2.**
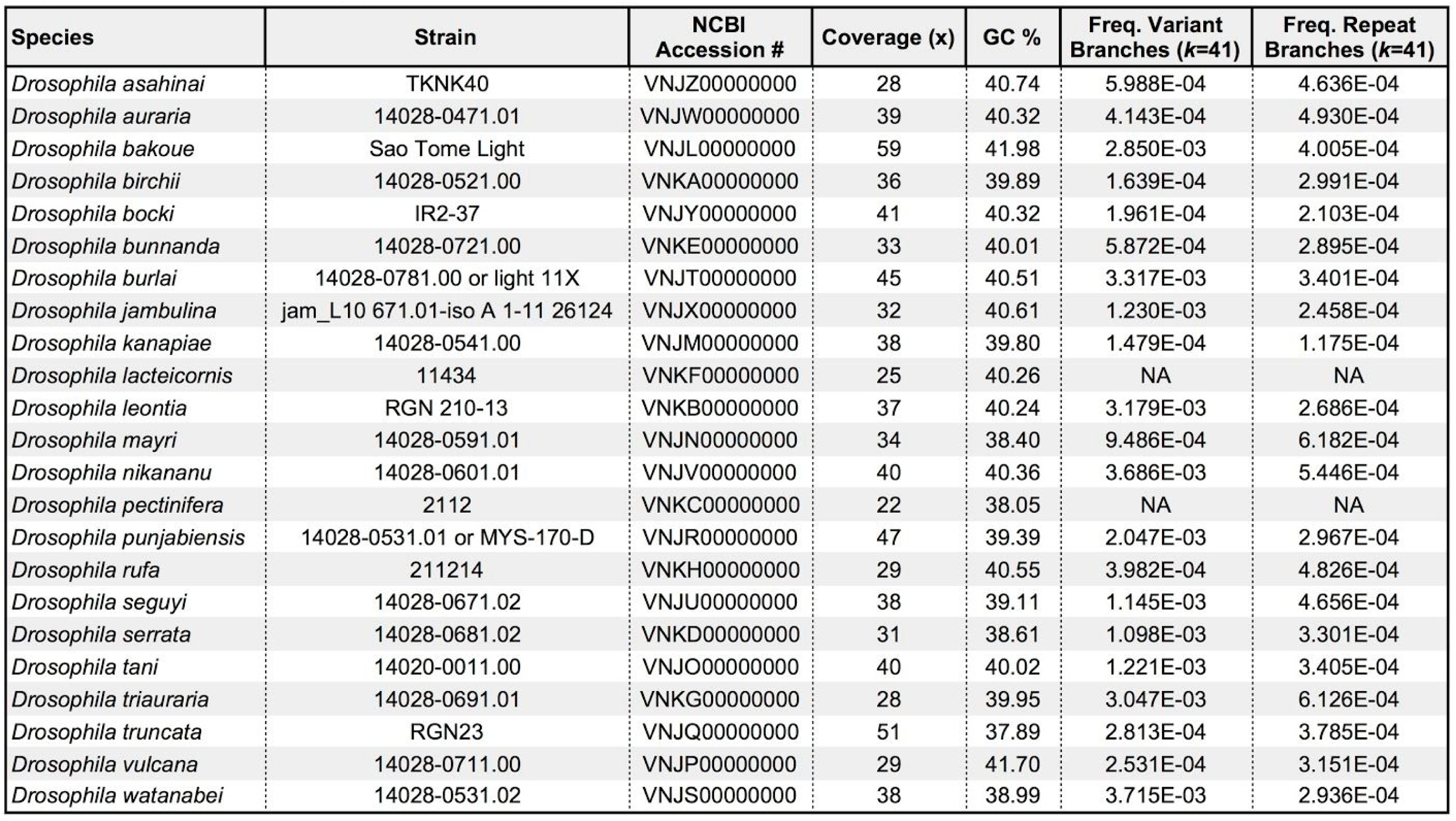
Additional sample and genome information. For *D. burlai* and *D. punjabiensis*, we sequenced one of two potential strains. Additional sequencing is underway to confirm the strain identification. Coverage is equal to the total amount of sequencing data (after read decontamination) divided by the estimated genome size (from SGA Preqc [21]). The GC % is based on the unassembled reads, not the assembly. See the Materials and Methods for additional information. The frequency of variant and repeat branches in the de Bruijn graph (*k*=41) was calculated by SGA Preqc. A *k*-mer size of 41 was chosen to maximize the number of species that could be compared. Sequence coverage was too low to estimate these parameters at *k*=41 for *D. lacteicornis* and *D. pectinifera*.

Estimated genome sizes within the *montium* group range from 155.1 Mb to 223.4 Mb (mean=196.4 Mb; median=198.1 Mb). These sizes are consistent with the previously assembled *D. kikkawai* [3] *and D. serrata* [19] genomes, with total sequence lengths of 164.3 Mb and 198.0 Mb, respectively. Our own *D. serrata* assembly (strain 14028-0681.02) has an estimated genome size of 184.7 Mb. The relatively small difference between our genome size estimate and the total contig length of the previously published PacBio *D. serrata* assembly (strain Fors4) [19] is likely a product of the imprecision of *k*-mer frequency spectrum-based genome size estimates, along with strain-level differences in genome size. Across all *montium* species, estimated genome size is strongly positively correlated with repeat content (*r*=0.88, *p*<1e-06).

Scaffold NG50s vary widely, from the remarkably contiguous *D. kanapiae* assembly (389,587 bp), to the highly fragmented *D. triauraria* assembly (17,513 bp). The contiguity of the *D. kanapiae* assembly is somewhat surprising (given the use of a small-insert library), but is related to genome and sample characteristics described below. The average scaffold NG50 across all *montium* species is 73,813 bp (median=54,224 bp).

Multiple factors can influence the contiguity of an assembly, including repeat content, heterozygosity, and sequencing depth. Large, repeat-rich genomes are typically difficult to assemble, as are highly heterozygous samples [21]. Given that the *montium* genomes were assembled using small-insert libraries (350 bp), they are especially sensitive to repeat content and heterozygosity. In the absence of long-distance information, in the form of mate-pair libraries or long-reads, large repeats form unresolvable structures in the graph. This results in fragmented assemblies that are missing many repeat copies [24,25]. Similarly, high levels of heterozygosity can create complicated graph structures that cause breaks in the assembly [26,27]. (Assemblers like Meraculous-2D [28] and Platanus [29] that are designed to handle high levels of heterozygosity typically require mate-pair libraries.) Finally, areas of low sequence coverage can also fragment an assembly [21].

Repeat content and heterozygosity vary widely across genomes / samples (Table 2), which in turn drive the scaffold NG50. For example, the *D. kanapiae* assembly owes its impressive contiguity to the lowest repeat content and heterozygosity level of any *montium* species. In contrast, the highly fragmented *D. triauraria* assembly combines the second highest repeat content with the fifth highest level of heterozygosity. To investigate the combined effect of repeat content, heterozygosity, and coverage on the scaffold NG50 across all *montium* species, we constructed a simple regression model (Table S1). As expected, the scaffold NG50 is inversely proportional to repeat content and heterozygosity, with repeat content impacting assemblies nearly twice as much as heterozygosity. The scaffold NG50 is generally unaffected by sequencing depth, as most genomes reach the minimum coverage necessary to effectively assemble contigs. While the sample size is small, the regression results are reassuring in that for each variable, the direction and relative magnitude of change is consistent with general genome assembly predictions.

The total scaffold length for most *montium* assemblies reaches 85 - 95 % of the estimated genome size. In Figure S1, this is where the curves intersect the x-axis. Given that our assemblies are missing many large repeat copies (see above), they should generally be shorter than the estimated genome size, with the magnitude of the difference proportional to the number, size, and divergence of repeat copies [30]. For example, the *D. pectinifera* and *D. mayri* assemblies reach only 67.8 % and 75.1 % of their estimated genome sizes, respectively. *D. mayri* has the highest repeat content of any *montium* species (Table 2), and *D. pectinifera* has the second largest estimated genome size (Table 1), which is highly correlated with repeat content. (The *D. pectinifera* sample was also heavily contaminated with bacteria. While the bacterial reads were filtered prior to assembly, their initial presence lowered the sequencing coverage of the fly genome. This further shortened the assembly, and prevented SGA Preqc [21] from estimating repeat content and heterozygosity.) In contrast, the relatively small and repeat-poor *D. kanapiae* genome yielded an assembly that reaches 97.9 % of its estimated genome size.

Compared to repeat content, heterozygosity can act as an opposing force on the total scaffold length. Given modest levels of heterozygosity, most assemblers collapse allelic variation into a single consensus sequence. As heterozygosity increases though, divergent haplotypes can sometimes be assembled independently on different scaffolds [26,27]. This artificially inflates the total scaffold length, and closes the gap between the estimated genome size and assembly length. (Some assemblers can also over-assemble the data and produce many small contigs / scaffolds known as “chaff” [31].) Consistent with this effect, the total scaffold lengths for *D. leontia* and *D. watanabei* actually exceed their estimated genome sizes. In Figure S1, these curves never intersect the x-axis. In the case of *D. watanabei*, this difference is large: 14.6 Mb. *D. watanabei* has the highest heterozygosity level of any *montium* species (Table 2), while *D. leontia* ranks fourth.

To investigate the combined effect of repeat content, heterozygosity, and coverage on the percentage of the estimated genome size that was assembled across all *montium* species, we constructed a simple regression model (Table S2). As expected, the percentage of the estimated genome size that was assembled is inversely proportional to repeat content, but positively correlated with heterozygosity - with repeat content being the primary driver. Once again, the sample size is small for a regression analysis, and the heterozygosity results only reach statistical significance at an alpha level of 0.10, but the results are generally as expected.

Overall, the *montium* assemblies are fragmented, as evidenced by their modest scaffold NG50s. However, taken in isolation, the NG50s say little about the quality of the assemblies. Any single metric (especially the NG50) can be a poor predictor of the quality / utility of an assembly. It is best to evaluate assemblies using a variety of methods, with an eye towards the downstream application [23]. For example, it is often advantageous to sacrifice contiguity for accuracy, and many questions can be answered without knowing the detailed repeat structure of the genome. We turn now to evaluating the *montium* assemblies in ways that will tell us if they are of sufficient contiguity and quality to study genes and transcriptional enhancers genome-wide.

### The vast majority of *montium* scaffolds are at least gene-sized

To study genes, a genome assembly should be present in at least gene-sized fragments [23,32]. By extension, such an assembly would also be useful for studying any features that are gene-sized or smaller, such as enhancers. Based on existing annotations of the PacBio *D. serrata* genome, the average gene length is up to 6.3 kb [19,33]. Figure 1 shows the relationship between the scaffold NG50 and the percentage of the assembly (total scaffold length) present in scaffolds that are at least 6.3 kb in length. Most *montium* assemblies are significantly shorter than their estimated genome sizes, on account of missing repeats. Therefore, we think it’s reasonable to ask the question: What percentage of the non-repetitive genome is present in at least gene-sized scaffolds? If we instead used the estimated genome sizes, the percentages would obviously decrease. Despite large differences in contiguity, all assemblies are present predominantly as scaffolds that are at least gene-sized. While there is a clear downward trend with decreasing NG50 (*r*=0.79, *p*<1e-5), in practice, this effect is modest. Even for the most fragmented assemblies, roughly 80 % of the assembly is present in at least gene-sized fragments.

**Figure 1.**
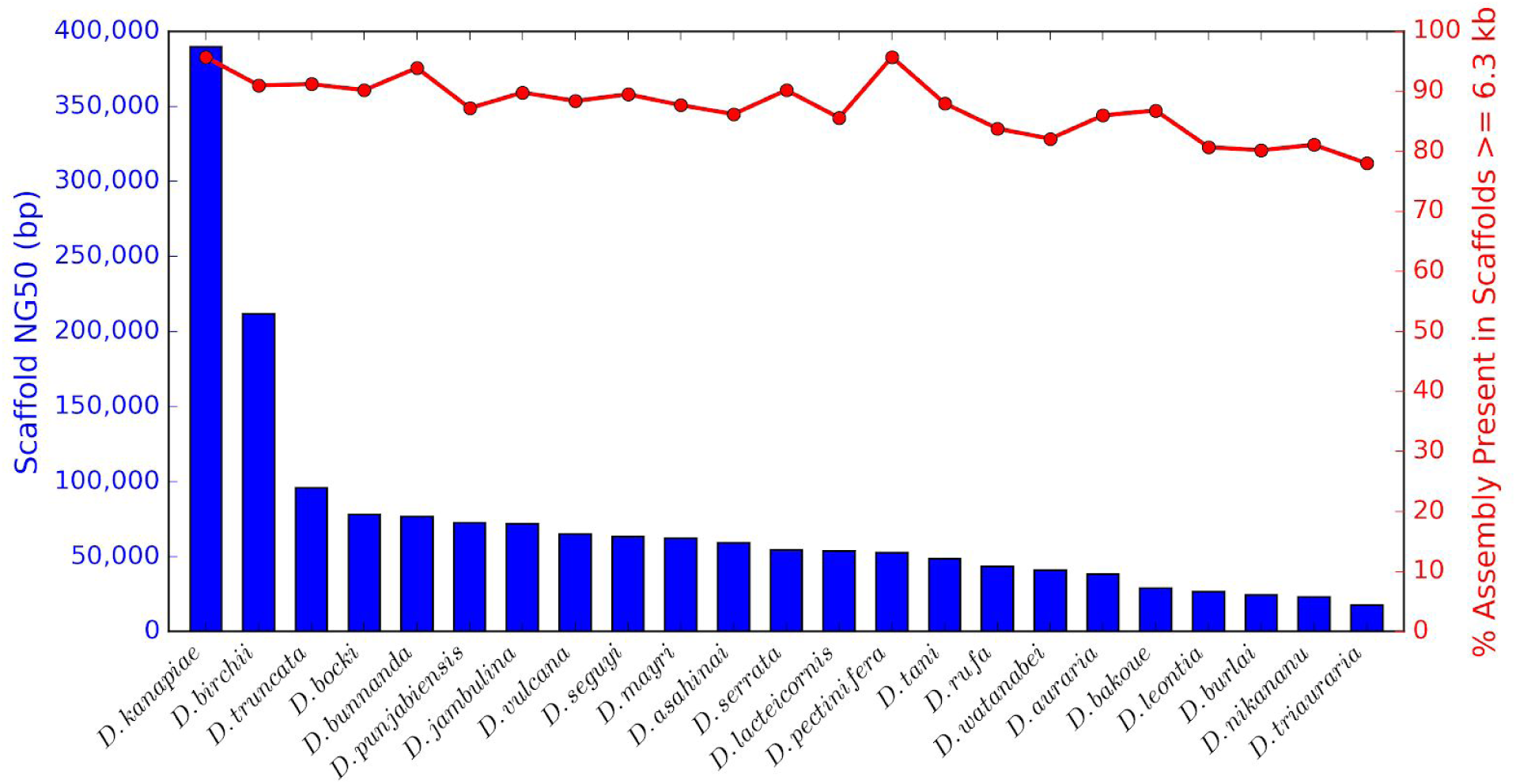
For all *montium* species, the vast majority of the assembly is present in at least gene-sized scaffolds, despite large differences in contiguity. Based on annotations of the previously assembled *D. serrata* genome [19,33], the average gene length is up to 6.3 kb. For each *montium* species, the blue bar graph shows the scaffold NG50, and the red line graph shows the percentage of the assembly (total scaffold length) present in scaffolds that are at least 6.3 kb in length. Species are listed in decreasing order of the scaffold NG50.

### All *montium* assemblies contain high percentages of known genes

The vast majority of scaffolds in each *montium* assembly are large enough to contain genes. However, do the scaffolds actually contain known genes? One way to assess the quality of an assembly is by annotation completeness: a good assembly should contain a high percentage of known genes. Benchmarking Universal Single-Copy Orthologs (BUSCOs) are single-copy genes present in more than 90 % of surveyed species [34,35]. The Dipteran BUSCO set contains 2,799 genes, and is based on a survey of 25 species. Figure 2 shows the BUSCO assessment results for eight *montium* assemblies. These species were chosen for their diversity: they occupy most subgroups in the *montium* group phylogeny [7]; include assemblies that fall far short of their estimated genome size; and represent a diversity of sample characteristics (heterozygosity and repeat content), genome size estimates, and assembly contiguity.

**Figure 2.**
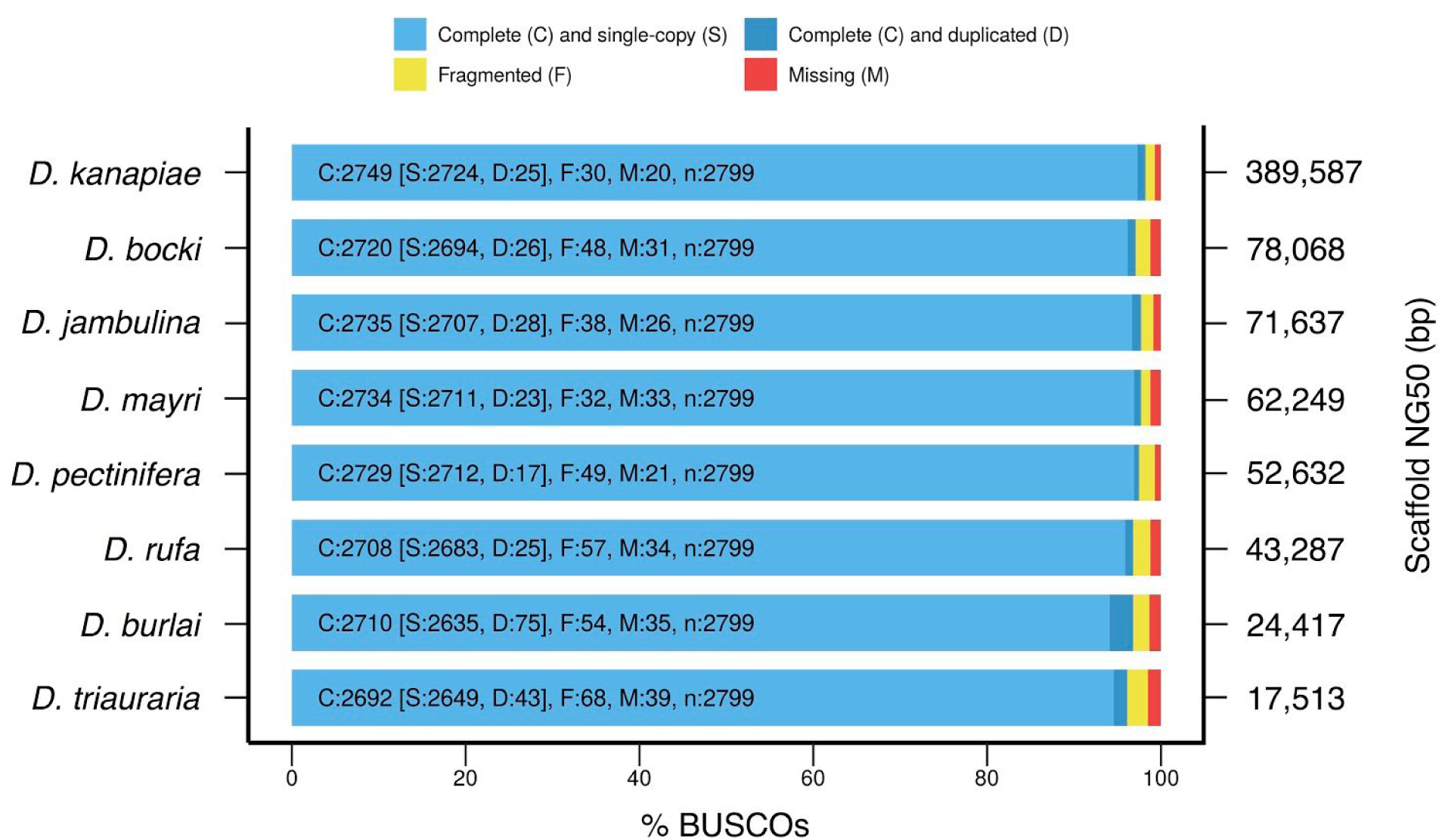
All *montium* assemblies contain high percentages of known genes despite large differences in contiguity. BUSCO [34,35] assessment results for eight *montium* genomes representing a diversity of genomes / assemblies. The Dipteran BUSCO set contains 2,799 genes. For each assembly, the bar graph reports the number of BUSCOs that are complete and single-copy, complete and duplicated, fragmented, and missing. The scaffold NG50 for each assembly is shown on the right.

They range from the small, repeat-poor, homozygous, and contiguous *D. kanapiae* (estimated genome size=155 Mb, scaffold NG50=390 kb), to the large, repeat-rich, highly heterozygous, and fragmented *D. triauraria* (estimated genome size=217 Mb, scaffold NG50=18 kb). Strikingly, despite the wide range of contiguities, there is little variation in gene content: at least 96.1 % of BUSCOs are complete (single-copy or duplicated) across all species. The *D. kanapiae* assembly exceeds 98 %. Ten BUSCOs are missing across all eight species, and likely represent lineage-specific loss events within Diptera. For comparison, the previously assembled *D. kikkawai* and *D. serrata* genomes, which approach scaffold / contig N50s of 1 Mb, reach 98.1 % and 96.2 %, respectively [19]. Once again, despite their relatively modest scaffold NG50s, our assemblies have performed well in metrics that matter for downstream analyses.

### Whole genome alignments of *montium* species to *D. melanogaster*

Given that the *montium* assemblies contain high percentages of known genes, we next determined if they also contain large percentages of known enhancer sequences. Non-coding regions are generally more difficult to assemble than genic regions. To facilitate the identification of enhancer sequences in *montium* genomes, we aligned each *montium* assembly to the *D. melanogaster* genome using a previously described whole genome alignment pipeline [36–38]. See the Materials and Methods for a complete description. Briefly, each *montium* assembly was individually aligned to the *D. melanogaster* genome using LASTZ [39]. The LASTZ alignments were then processed into structures called “chains” and “nets” [40] using a series of programs described in detail by [36]. The pipeline ultimately produced liftOver chain files. Given a set of coordinates for an annotated feature in the *D. melanogaster* genome, the liftOver [41] utility returns coordinates for the (putatively) orthologous sequence in an aligned *montium* genome. For this analysis, we also included the previously assembled *D. kikkawai* genome [3].

### All *montium* assemblies contain thousands of *D. melanogaster* enhancer orthologs

With the genomes aligned, we turned to looking for known enhancer sequences in the *montium* assemblies. We used a previously described set of 3,500 experimentally verified transcriptional enhancers that drive expression in the *D. melanogaster* embryo [42]. Using liftOver [41], we remapped the *melanogaster* coordinates onto each *montium* assembly. Across all *montium* assemblies, at least 99.6 % of enhancer coordinates were successfully remapped (Table 3). To determine whether the remapped coordinates correspond to orthologous sequence, we used BLASTn [43] to align the *montium* sequences back to the *melanogaster* genome, and the *melanogaster* sequences to the *montium* genomes. On average, 96.5 % of remapped coordinates are reciprocal best hits between the two genomes (Table 3). Of note, the highly contiguous *D. kikkawai* genome [3] is indistinguishable from our more fragmented assemblies. Next, we aligned each melanogaster sequence to its putative *montium* ortholog using BLASTn. Figure 3 shows illustrative results for the *D. lacteicornis* assembly, which is close to the median scaffold NG50. On average, 65.3 % of the *D. melanogaster* sequence aligns to sequence from *D. lacteicornis* (query coverage). The average percent identity is 75.1 %, and the Expect value (E) for the vast majority of alignments is essentially zero. Based on these results, it is clear that we can remap coordinates for thousands of *D. melanogaster* enhancers onto any *montium* assembly, and with a high level of confidence extract orthologous sequences.

**Table 3.** Thousands of orthologous *montium* enhancers can be identified by remapping *D. melanogaster* enhancer coordinates onto *montium* assemblies.

| Species | Attempted Remappings | Successful Remappings | % Successful Remappings | Reciprocal Best Hits (RBH) | % Successful Remappings that are RBH |
| --- | --- | --- | --- | --- | --- |
| *D. asahinai* | 3,457 | 3,450 | 99.8 | 3,361 | 97.4 |
| *D. auraria* | 3,457 | 3,448 | 99.7 | 3,347 | 97.1 |
| *D. bakoue* | 3,457 | 3,451 | 99.8 | 3,275 | 94.9 |
| *D. birchii* | 3,457 | 3,449 | 99.8 | 3,385 | 98.1 |
| *D. bocki* | 3,457 | 3,449 | 99.8 | 3,377 | 97.9 |
| *D. bunnanda* | 3,457 | 3,447 | 99.7 | 3,359 | 97.4 |
| *D. burlai* | 3,457 | 3,450 | 99.8 | 3,272 | 94.8 |
| *D. jambulina* | 3,457 | 3,450 | 99.8 | 3,327 | 96.4 |
| *D. kanapiae* | 3,457 | 3,449 | 99.8 | 3,406 | 98.8 |
| *D. kikkawai* | 3,457 | 3,449 | 99.8 | 3,377 | 97.9 |
| *D. lacteicornis* | 3,457 | 3,451 | 99.8 | 3,375 | 97.8 |
| *D. leontia* | 3,457 | 3,444 | 99.6 | 3,247 | 94.3 |
| *D. mayri* | 3,457 | 3,449 | 99.8 | 3,384 | 98.1 |
| *D. nikananu* | 3,457 | 3,451 | 99.8 | 3,221 | 93.3 |
| *D. pectinifera* | 3,457 | 3,449 | 99.8 | 3,383 | 98.1 |
| *D. punjabiensis* | 3,457 | 3,449 | 99.8 | 3,266 | 94.7 |
| *D. rufa* | 3,457 | 3,452 | 99.9 | 3,368 | 97.6 |
| *D. seguyi* | 3,457 | 3,444 | 99.6 | 3,334 | 96.8 |
| *D. serrata* | 3,457 | 3,447 | 99.7 | 3,350 | 97.2 |
| *D. tani* | 3,457 | 3,451 | 99.8 | 3,301 | 95.7 |
| *D. triauraria* | 3,457 | 3,448 | 99.7 | 3,258 | 94.5 |
| *D. truncata* | 3,457 | 3,449 | 99.8 | 3,366 | 97.6 |
| *D. vulcana* | 3,457 | 3,448 | 99.7 | 3,358 | 97.4 |
| *D. watanabei* | 3,457 | 3,449 | 99.8 | 3,147 | 91.2 |

Coordinates for *D. melanogaster* enhancers from [42] were remapped onto aligned *montium* assemblies using liftOver [41]. Reciprocal best hits (RBH) were identified by aligning *montium* sequences back to the *melanogaster* genome, and *melanogaster* sequences to the *montium* genomes - both using BLASTn [43]. See Materials and Methods for additional details. For comparison, we also included the previously assembled *D. kikkawai* genome [3].

**Figure 3.**
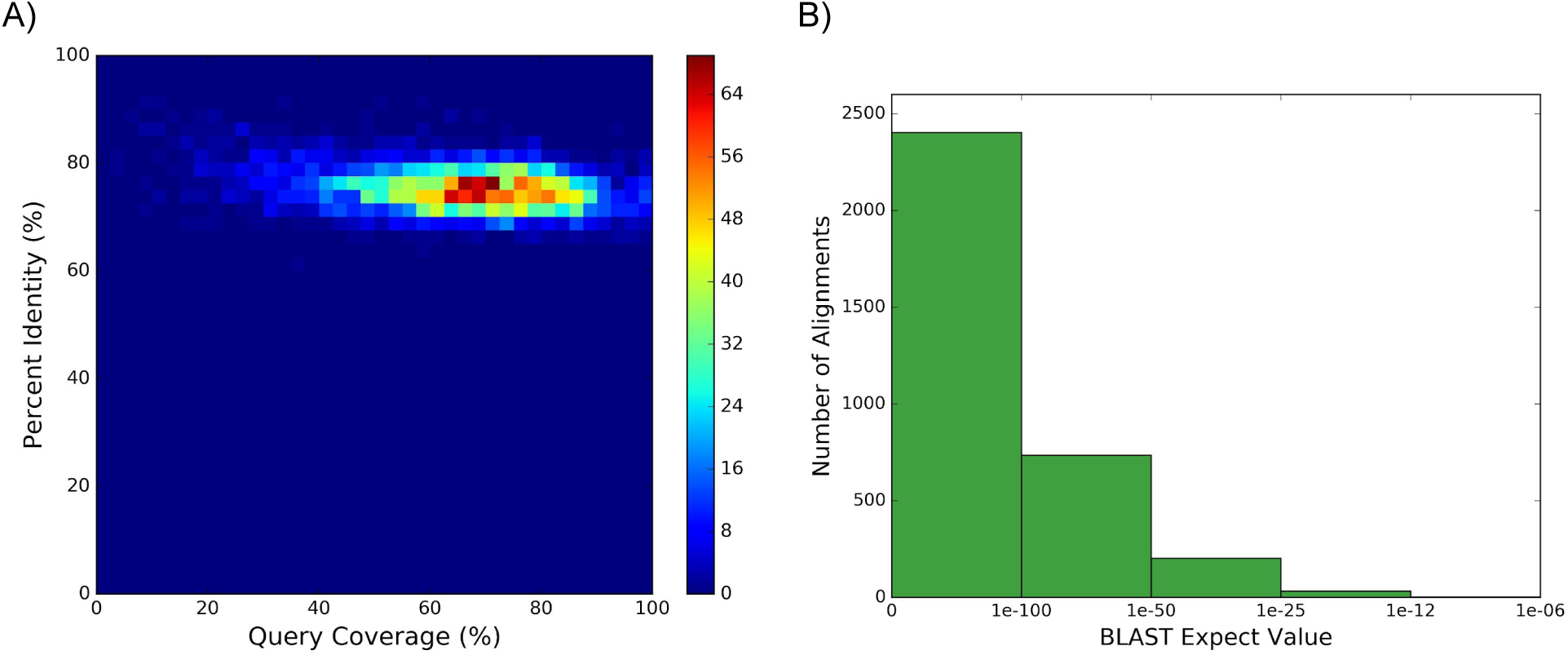
Pairwise BLASTn alignments between *D. melanogaster* enhancers and *D. lacteicornis* orthologs show highly similar sequences. 3,457 experimentally verified *D. melanogaster* enhancers from [42] *were remapped onto the D. lacteicornis* assembly using liftOver [41]. This yielded 3,375 reciprocal best hits between the *D. melanogaster* and *D. lacteicornis* genomes. *D. lacteicornis* was chosen for illustrative purposes because the assembly is close to the median scaffold NG50. A) 2D histogram showing query coverage and percent identity for 3,375 pairwise *D. melanogaster* - *D. lacteicornis* BLASTn [43] alignments. Query coverage is the percentage of *D. melanogaster* sequence that is aligned to *D. lacteicornis* sequence. B) Distribution of Expect values (E) for alignments in Part A.

### Identifying potential misassemblies

To look for large-scale misassemblies, we aligned the five longest scaffolds (up to 1 Mb) from our *D. serrata* assembly (strain 14028-0681.02) to orthologous contigs in the previously published - and far more contiguous - PacBio *D. serrata* assembly (strain Fors4) [19]. Absent large-scale misassemblies (e.g., translocations, relocations, and inversions), our scaffolds should generally align end-to-end within the longer PacBio contigs, with only relatively small insertions or deletions. Dotplots for pairwise alignments are shown in Figure 4. The first four alignments are highly collinear, with our scaffolds aligning end-to-end with only relatively small insertions / deletions. The fifth alignment is also highly collinear, but our scaffold (scf7180000629414) aligns across the ends of two PacBio contigs. The dotplot pattern also suggests the presence of inverted repeats in the vicinity of the breakpoint between contigs. To determine if this represents a potential misassembly, we next aligned scf7180000629414 to the orthologous scaffold in the previously published *D. kikkawai* assembly [3] (Figure S2). The alignment is once again highly collinear, but this time, our entire scaffold aligns end-to-end within the longer *D. kikkawai* scaffold. Unless the *D. kikkawai* scaffold is similarly misassembled, this indicates the overall structure of our scaffold is correct. However, the fact that scf7180000629414 spans a breakpoint in a PacBio assembly suggests either the repeat structure at this locus is more complicated in strain Fors4 than strain 14028-0681.02, or our scaffold contains a local repeat-induced misassembly.

**Figure 4.**
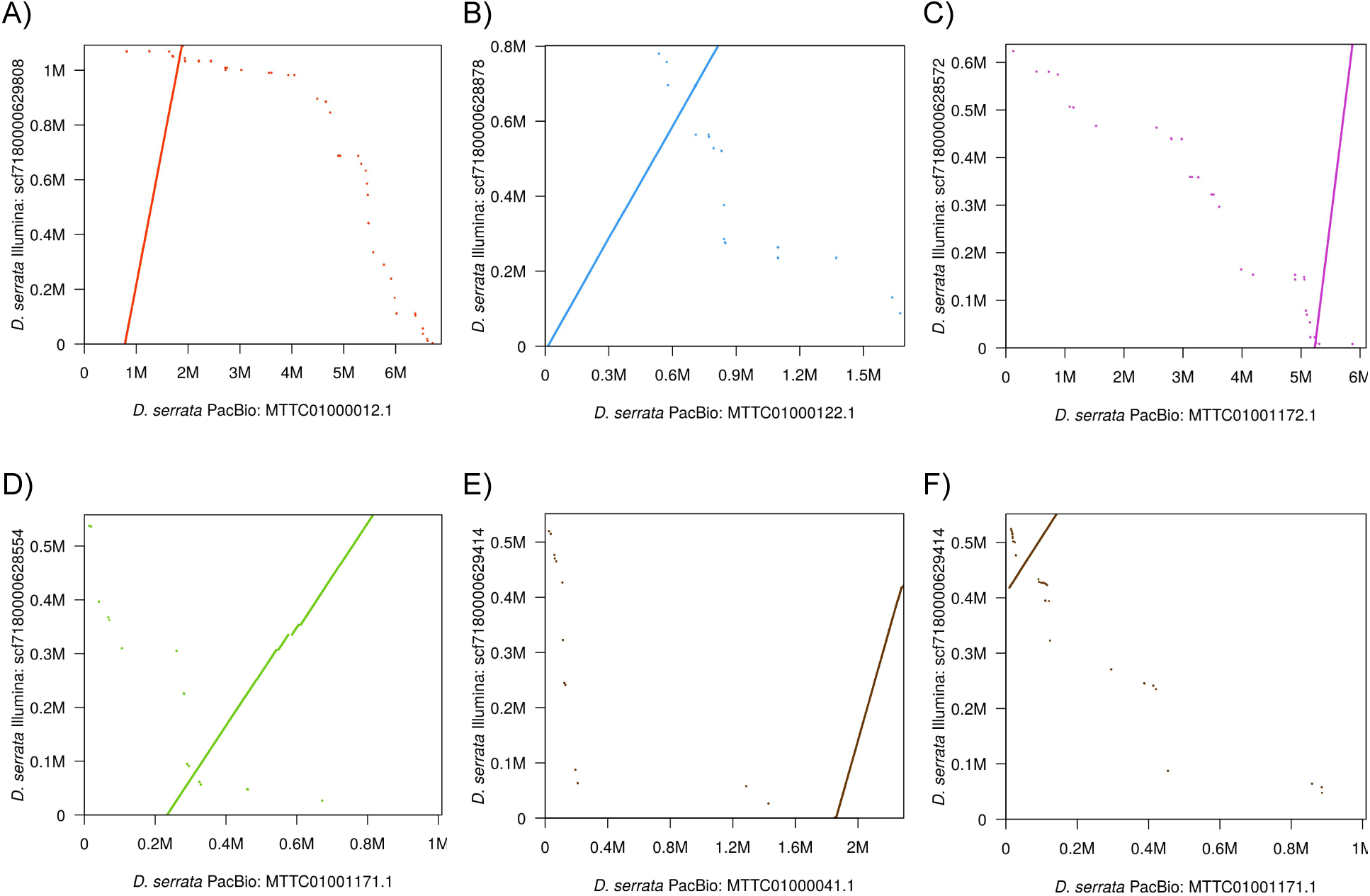
Alignments between the five longest scaffolds from our *D. serrata* assembly and orthologous contigs from a PacBio *D. serrata* assembly are highly collinear. Each dotplot shows the alignment of a scaffold from our Illumina *D. serrata* assembly (strain 14028-0681.02) to the orthologous contig from the previously published PacBio *D. serrata* assembly (strain Fors4) [19]. Pairwise alignments were generated by LASTZ [39]. Parts A) through D) show alignments for different scaffolds. Parts E) and F) show the alignment of the same scaffold to different contigs. Alignments are shown in decreasing order of scaffold length.

All draft genomes contain misassemblies, and ours are no different. While the above analysis generated reassuring results, it does not preclude the presence of other misassemblies (e.g., collapsed repeats, small inversions, or tandem alleles) within these scaffolds. We used REAPR [44] and Pilon [45] in our post-assembly pipeline to identify and correct as many errors as possible. While these programs work best with large-insert libraries (which we didn’t have), they nevertheless made significant improvements. We also “phased” our assemblies so that at each locus, the assembly represents the majority haplotype, within the limits of a small-insert library.

## Conclusions

We described the creation of a comparative genomic resource consisting of 23 genomes from the *Drosophila montium* species group, a large group of closely related species. Genomes for 22 of these species were presented here for the first time.

To make this endeavor financially feasible, we sequenced a single, small-insert library for each species. The absence of long-distance information made the assemblies especially sensitive to repeats and high levels of heterozygosity. As a result, many of the assemblies are fragmented, and the scaffold NG50s vary widely based on genome / sample characteristics. The total scaffold length of most assemblies is also significantly shorter than the estimated genome sizes.

However, just because most assemblies are fragmented, does not mean they are poor quality. Quite to the contrary, the BUSCO [34,35] analysis showed that all assemblies, regardless of contiguity, contain at least 96 % of known single-copy Dipteran genes (n=2,799). Similarly, by aligning our assemblies to the *D. melanogaster* genome and remapping coordinates for a large set of enhancers (n=3,457) [42], we showed that each *montium* assembly contains orthologs for at least 91 % of *D. melanogaster* enhancers. (This same approach can be used for any annotated feature in the *D. melanogaster* genome.) Importantly, the genic and enhancer contents of our assemblies are comparable to that of far more contiguous *Drosophila* assemblies. Finally, the alignment of our *D. serrata* assembly to a previously published PacBio *D. serrata* assembly [19] showed that our longest scaffolds (up to 1 Mb) are free of large-scale misassemblies.

While all of our assemblies are complete enough to study genes and enhancers, if other researchers are interested in repeat structure, any *montium* assembly can be improved on an as-needed basis. By pairing our short-read data (all of which is publicly available) with mate-pair libraries or PacBio long-reads, they can easily generate vastly more contiguous assemblies that include most repeat copies.

Going forward, our genome assemblies will be a valuable resource that can be used to further resolve the *montium* group phylogeny; study the evolution of protein-coding genes and enhancers; and determine the genetic basis of ecological and behavioral adaptations.

## Materials and Methods

### Fly lines

Fly lines for each *montium* species listed in Table 2 were gifts of Artyom Kopp and Michael Turelli, or were acquired from the *Drosophila* Species Stock Center. Additional strain information can be found in the associated BioSample record maintained by NCBI (see Data Availability below).

All fly lines were maintained in small population vials. Prior to sequencing, some lines went through several generations of inbreeding. Other lines were not inbred, either due to difficulty maintaining the fly line, or time limitations.

### Library preparation and sequencing

For each species, DNA was extracted from three female flies using the QIAGEN QIAamp DNA Micro Kit. Sequencing libraries were constructed using the Illumina TruSeq DNA PCR-Free Kit for 350 bp inserts, and visualized on Agilent High Sensitivity DNA chips. Libraries were clustered on Illumina HiSeq 2000 or HiSeq 2500 Systems, generating 100 bp paired-end reads. All sequencing was done at the Vincent J. Coates Genomics Sequencing Laboratory at UC Berkeley. Multiple species were pooled on each lane in an effort to reach sequencing depths of at least 30x per species, assuming genome sizes around 164 Mb (based on the previously published *D. kikkawai* genome [3]).

### Read exploration and pre-processing

Prior to assembly, read quality and genome / sample characteristics (e.g., estimated genome size, repeat content, and heterozygosity) were explored using FastQC (v. 0.11.2) [46] and String Graph Assembler (SGA) Preqc (v. 0.10.15) [21]. SGA Preqc was run using the following commands: sga preprocess, with the option --pe-mode 1; sga index, with the options -a ropebwt and --no-reverse; and sga preqc. The report was generated using the included script sga-preqc-report.py.

Reads from some sequencing runs contained an extra base (i.e., 101 bases instead of 100). This extra base was trimmed using BBDuk (BBMap v. 36.11) [47], with the option ftr=99. Reads were adapter-trimmed for known Illumina adapters using BBDuk, with the options ktrim=r, k=23, mink=9, hdist=1, minlength=75, tpe=t, and tbo=t. The adapter-trimmed reads were then quality-trimmed to Q10 using BBDuk (which implements the Phred algorithm), with the options qtrim=rl, trimq=10, and minlength=51.

The *D. bakoue* library was sequenced across two lanes. Sequence quality on the first lane was adversely affected by problematic tiles, as evidenced by the Per Tile Sequence Quality plot generated by FastQC [46]. Low-quality reads were removed using FilterByTile (BBMap v. 37.56) [47], using a statistical profile that included other libraries on the same flowcell. To lower the total sequencing coverage from approximately 75x to 60x, filtered reads from the first lane were subsampled using Reformat (BBMap v. 36.11), with the option samplerate=0.6.

### Read decontamination

Sequence contaminants were identified by reviewing the Per Sequence GC Content plots from FastQC [46], and the GC Bias plots from SGA Preqc [21]. Contaminants formed secondary peaks or spikes in the Per Sequence GC Content plots, and secondary GC % - *k*-mer coverage clusters in the GC Bias plots.

The *D. pectinifera* and *D. vulcana* sequencing libraries were heavily contaminated with microorganisms (mostly bacteria). Low levels of bacteria were also present in the *D. burlai* library. We utilized two different decontamination strategies.

For *D. pectinifera*, we adopted a decontamination strategy similar to [48]. The reads were first assembled using SOAPdenovo2 [49], with the options -K 49 and -R. Assembled scaffolds at least 1 kb in length were used to create a GC % vs. average k-mer coverage plot. Scaffolds with 35 <= GC % <= 66 and 40.5 <= average *k*-mer coverage <= 68 were classified as candidate contaminant scaffolds. To avoid removing *Drosophila* scaffolds, candidate contaminant scaffolds were aligned to sequences in NCBI’s Nucleotide database [50] using BLASTn (v. 2.2.31+) [43]. Candidate contaminant scaffolds that aligned to known microorganism sequences were used to create a contaminant reference. Finally, the original reads were aligned to the contaminant reference using Bowtie 2 (v. 2.2.3) [51], with the option --local, and pairs of reads that aligned concordantly were removed prior to the subsequent assembly.

For *D. vulcana* and *D. burlai*, 10,000 reads were sampled from the R1 FASTQ files using seqtk sample (v. 1.0-r75-dirty) [52], and then converted to FASTA format using seqtk seq. After reviewing the Per Sequence GC Content plot from FastQC [46], potential sequence contaminants were isolated based on their GC %, and then aligned to sequences in NCBI’s Nucleotide database [50] using BLASTn [43]. This led to the identification of closely related bacteria and yeast genomes, which were combined into a contaminant reference. Finally, the original reads were aligned to the contaminant reference using Bowtie 2 [51], with the option --local, and pairs of reads that aligned concordantly were removed prior to assembly.

The *D. burlai, D. jambulina, D. mayri, D. seguyi*, and *D. vulcana* libraries appeared to be contaminated with highly abundant individual sequences (or groups of similar sequences). These sequences created spikes in the Per Sequence GC Content plots from FastQC [46], and corresponded to eight-bp or ten-bp simple sequence repeats (SSRs) that were present in both the forward and reverse reads of the same DNA fragment. The origin of the sequences was unclear. Once the potential contaminant sequences were identified, matching sequences were removed from the reads using BBDuk [47], with the options k=75 and hdist=1.

### Genome GC %

The GC % for each species was calculated using the unassembled reads. Given that the assemblies are depleted of large repeat copies, we thought this approach would produce more accurate estimates than simply calculating the GC % of the assemblies. (That being said, raw sequencing data can also have GC biases.) Base frequency and read length histograms were constructed using the adapter-trimmed and decontaminated R1 FASTQ files and BBDuk [47], with the options bhist, lhist, and gcbins=auto. The output was then used to calculate the GC % of the reads, which are reported in Table 2. On average, the GC % of the unassembled reads is 1.5 % lower than the GC % of the assemblies (data not shown).

### Choosing an assembler

Exploration of the data using SGA Preqc [21] showed that the *montium* genomes / samples represent a diversity of genome size estimates, repeat contents, heterozygosity levels, and sequencing error rates. Extensive tests were conducted to identify the assembler that performed the best across these diverse samples.

We tested the following assemblers: ABySS [53], MaSuRCA [20], Meraculous-2D [28], SOAPdenovo2 [49], SPAdes [54] / dipSPAdes [55], and Velvet [56]. The resulting assemblies were evaluated using a number of metrics, including contiguity statistics, REAPR [44], Feature Response Curves (*FRC*^*bam*^) [57], BUSCO assessments [34,35], and the scrutiny of individual enhancer sequences.

### Primary assemblies

All genomes were assembled using MaSuRCA (v. 3.2.2) [20], on a server with 48 Intel Xeon E5-2697 v2 2.70 GHz processors and 377 GB of RAM. Assemblies could use up to 36 CPUs. MaSuRCA was supplied with reads that had been force-trimmed to 100 bp and decontaminated, but not adapter-trimmed or quality-trimmed. The authors of MaSuRCA recommend no read trimming, and MaSuRCA performs error correction internally using QuorUM [58].

The configuration file for each species contained the insert-size mean and standard deviation for the corresponding sequencing library, as well as the following parameters: GRAPH_KMER_SIZE=auto, USE_LINKING_MATES=1, CA_PARAMETERS=cgwErrorRate=0.15, KMER_COUNT_THRESHOLD=1, and SOAP_ASSEMBLY=0. The Jellyfish hash size (JF_SIZE) was set to the product of the estimated genome size and coverage.

### Post-assembly pipeline

Our post-assembly pipeline started with assemblies present in the MaSuRCA [20] output directory 9-terminator, so we could control the gap closing process.

For each assembly, MaSuRCA [20] created a small number of scaffolds with massive gaps (tens of kb in length). Given the insert-sizes of the sequencing libraries (∼ 350 bp), these gaps had to be erroneous. Therefore, scaffolds were split on any gap that was unreasonably large relative to the insert-size of the library. Maximum allowed gap sizes were typically around 200 - 600 bp, depending on the library.

REAPR - Recognition of Errors in Assemblies using Paired Reads (v. 1.0.18) [44] was used to identify errors in the assemblies, and to generate new “broken” assemblies that were split on errors occurring over gaps. Errors within contigs were hard-masked with Ns. The command reapr smaltmap was used to align adapter-trimmed reads to the assemblies, and reapr pipeline generated the broken assemblies. Sequences starting with “REAPR_bin” (i.e., the original unmasked sequence) were later filtered from the broken assemblies.

Gaps in the assemblies were closed using a two-step process with adapter-trimmed and quality-trimmed reads. The first round of gap closing was performed using GapCloser (v. 1.12) [49]. This also helped to identify tandem alleles (a type of misassembly) [59], which GapCloser left as single-N gaps. The second round was done using Sealer (abyss-sealer v. 2.0.2) [60], with the option -P 10. For each assembly, “*k* sweeps” typically ranged from *k*=80 to *k*=30 (in decrements of 10), but varied if Sealer became stuck on a given *k*-mer size. After two rounds of gap closing, the *D. triauraria* assembly contained more than 2,000 single-N gaps. The remaining single-N gaps (and associated flanking sequence) were hard-masked with 300 Ns, and Sealer was run a second time using the above settings. This potentially extended the flanking sequence extracted by Sealer beyond the boundaries of the original tandem allele, thereby making it possible to find a connecting path in the graph. This decreased the number of single-N gaps below 2,000.

The assemblies were further improved using Pilon (v. 1.22) [45], an automated variant detection and genome assembly improvement tool. Adapter-trimmed reads were first aligned to the assemblies using Bowtie 2 [51], with the option --very-sensitive-local. Pilon was then run with the options --fix all,amb, --diploid, and --mingap 1. This attempted to fix SNPs, indels, local misassemblies, and ambiguous bases, as well as fill remaining gaps.

After running Pilon [45], adapter-trimmed reads were aligned to the improved assemblies using Bowtie 2 [51], with the option --very-sensitive-local. Detailed inspection of the aligned reads showed that many scaffolds were mosaics of multiple haplotypes present in the original samples. This was a significant problem for highly heterozygous samples, as it created numerous recombinant haplotypes not present in the original samples. Our goal therefore was to create “phased” assemblies that reflected the majority haplotype at each variable locus.

Pilon [45] was run a second time on the improved assemblies, but this time it was used as a variant detection tool to generate VCF files (option --vcf). For highly heterozygous samples, multiple overlapping variants were sometimes present at the same locus, which often led to aberrant phasing behavior. Variants can overlap because they share the same start position, or a large deletion might overlap SNPs or smaller indels. The VCF files were filtered so that only one overlapping variant was retained: either the structural variant (if one was present), or the majority variant. Variants in the VCF files were phased using the read-based phasing tool WhatsHap (v. 0.14.1) [61], with the options phase, --ignore-read-groups, --tag=PS, and --indels. BCFtools (v. 1.5) [62] with the options view, --phased or --exclude-phased was then used to create VCF files with only phased or un-phased variants. To facilitate parsing of the phased VCF files, a sequence dictionary was first created with the tool CreateSequenceDictionary from Picard (v. 2.12.1-SNAPSHOT) [63], and then VariantsToTable from the Genome Analysis Toolkit (GATK) (v. nightly-2017-09-13-g315c945) [64] was used to create tab-delimited tables of variants. For each phase set in the table, the majority haplotype was determined based on the cumulative read count of variants on each haplotype (A or B), with indels weighted half as much as SNPs (because of alignment issues with indels). Phased variants that were present on majority haplotypes were retained. For un-phased variants, the majority allele was retained. New VCF files were then created using only the retained phased and un-phased variants. Finally, BCFtools consensus was used to create new “phased” assemblies by applying the variants in these VCF files to the original “un-phased” assemblies.

Lastly, any remaining ambiguous bases (except N) were randomly assigned to a single base, and scaffolds shorter than 1 kb in length were removed.

### Assembly decontamination

Contaminants in the final assemblies were identified by NCBI’s Contaminant Screen. Most assemblies contained small numbers of scaffolds from bacterial or yeast species, which were removed. Four scaffolds across all assemblies also contained suspected adapter / primer sequences. These scaffolds were split on the potential contaminant.

### Assembly statistics

The correlation between the estimate genome size and the log_10_(repeat content) was calculated using the R (v. 3.4.1) [65] function cor.test(), with the option method=“pearson”. Repeat content is the frequency of repeat branches in the de Bruijn graph (*k*=41), as calculated by SGA Preqc [21].

Regression models for predictors of scaffold NG50, and the percentage of the estimated genome size that was assembled, were constructed using the R [65] function lm().

### Percentage of assembly present in gene-sized scaffolds

Figure 1 style adapted from figure in [23]. The correlation between the log_10_(scaffold NG50) and the percentage of the assembly present in scaffolds greater than or equal to 6.3 kb in length was calculated using the R [65] function cor.test(), with the option method=“pearson”.

### BUSCO assessment

The assemblies were searched for known genes using BUSCO (v. 3.0.2) [34,35], with the profile library diptera_odb9. The following options were specified in the configuration file: mode=genome, evalue=1e-3, limit=3, and long=False. The BUSCO plot was constructed using the included script generate_plot.py.

### Whole genome alignment pipeline

*Each montium* genome was individually aligned to the *D. melanogaster* genome (NCBI Assembly ID: 202931, Release 6 plus ISO1 MT / UCSC Genome Browser Assembly ID: dm6) [66–68] using a previously described whole genome alignment pipeline [36–38]. Target and query genomes were soft-masked using RepeatMasker (v. open-4.0.7) [69], with the sequence search engine RMBlast (v. 2.2.28) and Tandem Repeat Finder (TRF) (v. 4.04) [70], and the options -s, -species drosophila, -gccalc,-nocut, and -xsmall. Pairs of genomes were aligned using LASTZ (v. 1.04.00) [39], with the following options from [3]: target_genome[multiple], --masking=50,--hspthresh=2200, --ydrop=3400, --gappedthresh=4000, --inner=2000, and --format=axt. The LASTZ alignments were then processed into structures called “chains” and “nets” [40] using a series of programs described in detail by [36]. Briefly, FASTA files for the target and query assemblies were converted to 2bit format using faToTwoBit. Files containing chromosome / scaffold lengths were created using faSize with the option-detailed. Gapless alignments (“blocks”) were linked together into maximally scoring chained alignments, or chains. The order of blocks within chains must be the same in both target and query genomes. Blocks within chains can be separated by insertions / deletions, inversions, duplications, or translocations. Chains were built using axtChain with the option -linearGap=medium, and then filtered using chainPreNet. Gaps in high-scoring chains were filled in with lower scoring chains, creating hierarchies (parent-child relationships) known as nets. Nets were constructed using chainNet with the option -minSpace=1, and then annotated using netSyntenic. Finally, subsets of chains found in nets were extracted using netChainSubset, creating liftOver chain files.

### Identification of *montium* sequences orthologous to *D. melanogaster* enhancers

[42] previously described a large set of DNA fragments (Vienna Tiles) that drive expression in the *D. melanogaster* embryo. A total of 3,457 fragments were positive for enhancer activity and PCR-verified. *D. melanogaster* coordinates were remapped onto each *montium* assembly using liftOver [41], with the options -minMatch=0.1 and-multiple. The liftOver program was originally written to remap coordinates between assemblies of the same species. However, it is routinely used for interspecies lifts, and in our experience, it performed well. In cases of multiple remappings for a single fragment, the larger coordinate span was retained, as it typically contained the sequence of interest.

The following strategy was used to identify reciprocal best hits. The candidate orthologs from each *montium* assembly were aligned back to the *D. melanogaster* genome using BLASTn [43], with the options -evalue 0.00029, -word_size 11, -reward 2, -penalty -3, -gapopen 5, -gapextend 2, -dust no, and -outfmt 6; and BEDTools (v. 2.17.0) [71] intersect was used to determine whether the highest scoring BLAST hit for each montium sequence overlapped the original fragment coordinates in the *melanogaster* genome. Conversely, the *melanogaster* fragment sequences were aligned to each *montium* assembly using BLASTn, and BEDTools intersect was used to determine whether the highest scoring BLAST hit for each *melanogaster* sequence overlapped the remapped fragment coordinates in the *montium* assembly. Fragments that met both criteria were classified as reciprocal best hits.

To visualize the similarity between reciprocal best hits, pairs of *montium* (subject) and *melanogaster* (query) sequences were aligned using BLASTn [43], with the options-task blastn-short, -evalue 0.00029, -reward 2, -dust no, and -outfmt 6. The BLAST output was filtered so that lower-scoring hits nested within, or partially overlapping, higher-scoring hits were removed / trimmed. The resulting hits were used to calculate the query coverage and length-weighted percent identity for the alignment. The lowest E value for each pairwise alignment was used for the 1D histogram.

### Scaffold alignment visualization using dotplots

We aligned our *D. serrata* assembly (strain 14028-0681.02) to the previously published *D. kikkawai* [3] and PacBio *D. serrata* (strain Fors4) [19] assemblies using the whole genome alignment pipeline detailed above [36–38].

Pairs of orthologous scaffolds / contigs were aligned for visualization using LASTZ [39], with the following options (in part from [3]): --chain, --masking=50,--hspthresh=2200, --ydrop=3400, --gappedthresh=4000, --inner=2000, and--format=rdotplot. For consistent visualization, our scaffolds scf7180000628572 and scf7180000629414 were reverse-complemented prior to pairwise alignment. Dotplots were constructed using R [65].

### Scripting and plotting

Unless otherwise stated, all scripts were written in Python (v. 2.7.14) [72], and plots were created using Matplotlib (v. 1.5.1) [73].

## Author Contributions

MJB assembled and analyzed all of the genomes, wrote the original draft of the paper, edited / revised the paper, and constructed the first 11 sequencing libraries. CCM and HAW constructed the remaining 12 sequencing libraries. CCM also edited / revised the paper. MBE conceived of the overall project, provided funding, supervised the project, and edited / revised the paper.

## Data Availability

All assemblies and sequencing data are publicly available through the *Drosophila montium* Species Group Genomes Project, NCBI BioProject Accession PRJNA554346. This record provides links to the assemblies, BioSamples, and sequencing data. The Whole Genome Shotgun projects have been deposited at DDBJ/ENA/GenBank under the accession numbers listed in Table 2. The versions described in this paper are versions XXXX01000000. Raw sequencing data was uploaded to the NCBI Sequence Read Archive (SRA). Besides removing reads that did not pass filtering, the FASTQ files were unprocessed.

Sequencing libraries for *D. bakoue, D. kanapiae, D. mayri, D. punjabiensis, D. tani, D. truncata*, and *D. vulcana* were spread across two lanes of a flowcell. When the FASTQ files were uploaded to the NCBI SRA, the R1 and R2 files from both lanes were combined into individual R1 and R2 FASTQ files. If users wish to demultiplex reads by lane for these samples, lane information (always 1 or 2) is preserved in the sequence identifier line of the original FASTQ files.

Soft-masked assemblies and liftOver chain files were deposited in the Dryad repository: https://doi.org/10.6078/D1CH5R.

## Acknowledgements

This work was supported by a Howard Hughes Medical Institute (HHMI) investigator award to MBE. The authors also wish to thank Michael Turelli, Artyom Kopp, Emily Delaney, Brandon Cooper, and Will Conner for their help working with *montium* species.

---

**Table S1.** Regression analysis for predictors of scaffold NG50.

| Predictor Variables | Model 1 | Model 2 |
| --- | --- | --- |
| Intercept | 0.822<br>(0.768) | 0.850<br>(0.745) |
| COVERAGE | 1.63e-03<br>(4.90e-03) |  |
| HETEROZYGOSITY | -0.420 ***<br>(8.69e-02) | -0.413 ***<br>(8.17e-02) |
| REPEAT CONTENT | -0.747 **<br>(0.242) | -0.763 **<br>(0.231) |
| Multiple $R^2$ | 0.76 | 0.76 |
| Adjusted $R^2$ | 0.72 | 0.73 |

N=21 for all models. Standard errors in parentheses.
Scaffold NG50 = $\log_{10}(\text{Scaffold NG50})$
COVERAGE = total sequenced bases (after decontamination) / estimated genome size
HETEROZYGOSITY = $\log_{10}(\text{frequency of variant branches in de Bruijn graph, } k=41)$
REPEAT CONTENT = $\log_{10}(\text{frequency of repeat branches in de Bruijn graph, } k=41)$
Estimated genome sizes and the frequency of variant / repeat branches were calculated by SGA Preqc [21].

**Table S2.** Regression analysis for predictors of the percentage of the estimated genome size that was assembled.

| Predictor Variables | Model 1 | Model 2 |
| --- | --- | --- |
| Intercept | 5.77<br>(25.5) | 4.65<br>(24.8) |
| COVERAGE | -6.50e-02<br>(0.163) |  |
| HETEROZYGOSITY | 5.43 .<br>(2.89) | 5.13 .<br>(2.72) |
| REPEAT CONTENT | -30.1 **<br>(8.04) | -29.4 **<br>(7.69) |
| Multiple $R^2$ | 0.46 | 0.45 |
| Adjusted $R^2$ | 0.36 | 0.39 |

N=21 for all models. Standard errors in parentheses.
% of est. genome size assembled = (assembly length / estimated genome size) \* 100
COVERAGE = total sequenced bases (after decontamination) / estimated genome size
HETEROZYGOSITY = $\log_{10}$ (frequency of variant branches in de Bruijn graph, $k=41$ )
REPEAT CONTENT = $\log_{10}$ (frequency of repeat branches in de Bruijn graph, $k=41$ )
Estimated genome sizes and the frequency of variant / repeat branches were calculated by SGA Preqc [21].

**Figure S1.**
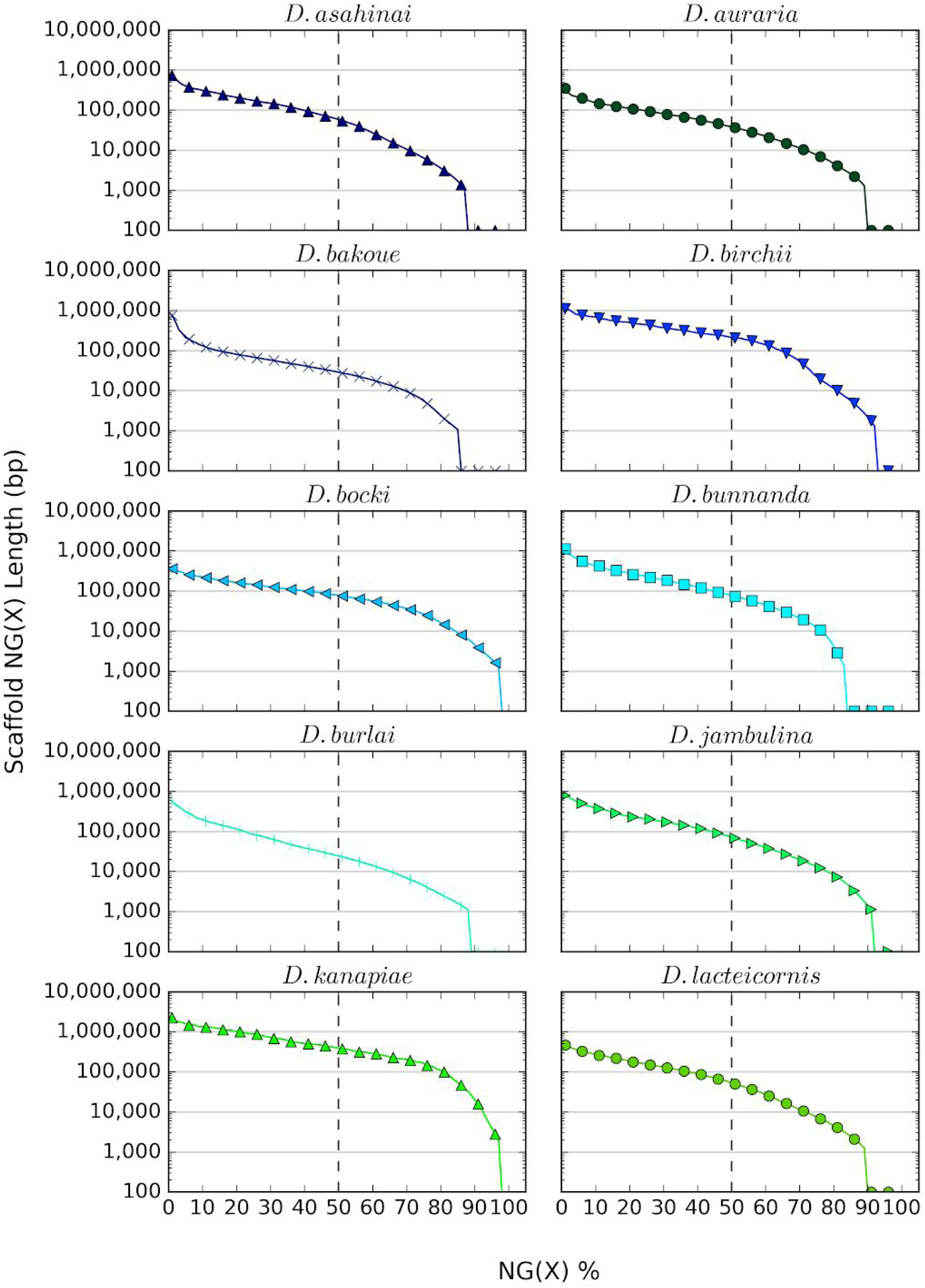
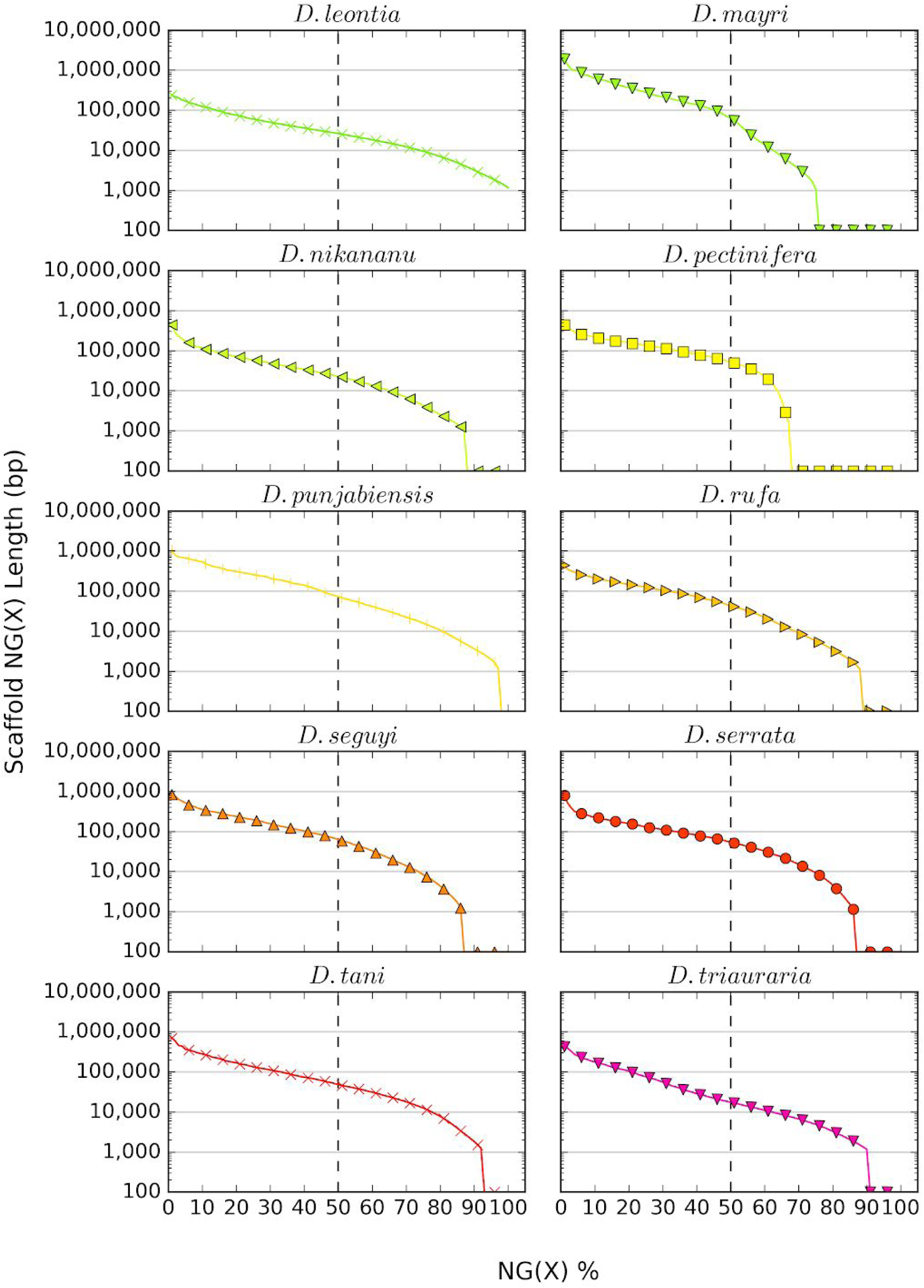
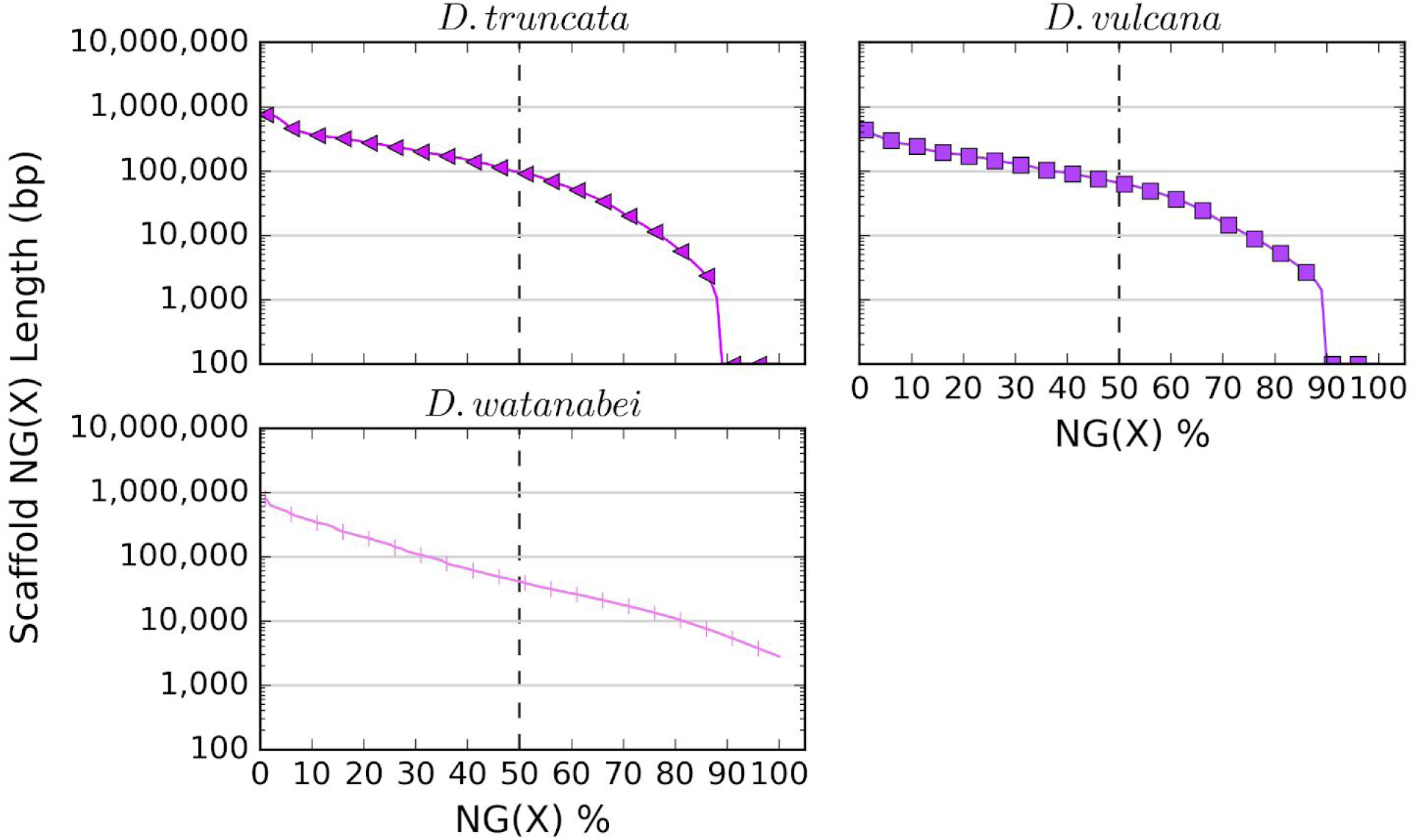
NG graphs showing the distribution of scaffold lengths for 23 *montium* assemblies. To calculate the scaffold NG50 [22,23], scaffold lengths are ordered from longest to shortest and then summed, starting with the longest scaffold. The NG50 is the scaffold length that brings the sum above 50 % of the estimated genome size. When this calculation is repeated for all integers from 1 to 100, the result is an NG graph [23]. NG graphs were constructed for each *montium* species using the corresponding genome size estimates from SGA Preqc [21]. When a series intersects the x-axis, it means the total scaffold length is shorter than the estimated genome size. Similarly, if the series never touches the x-axis, then the assembly is longer than the estimated genome size. Due to filtering, the shortest scaffold present in any assembly is 1 kb.

**Figure S2.**
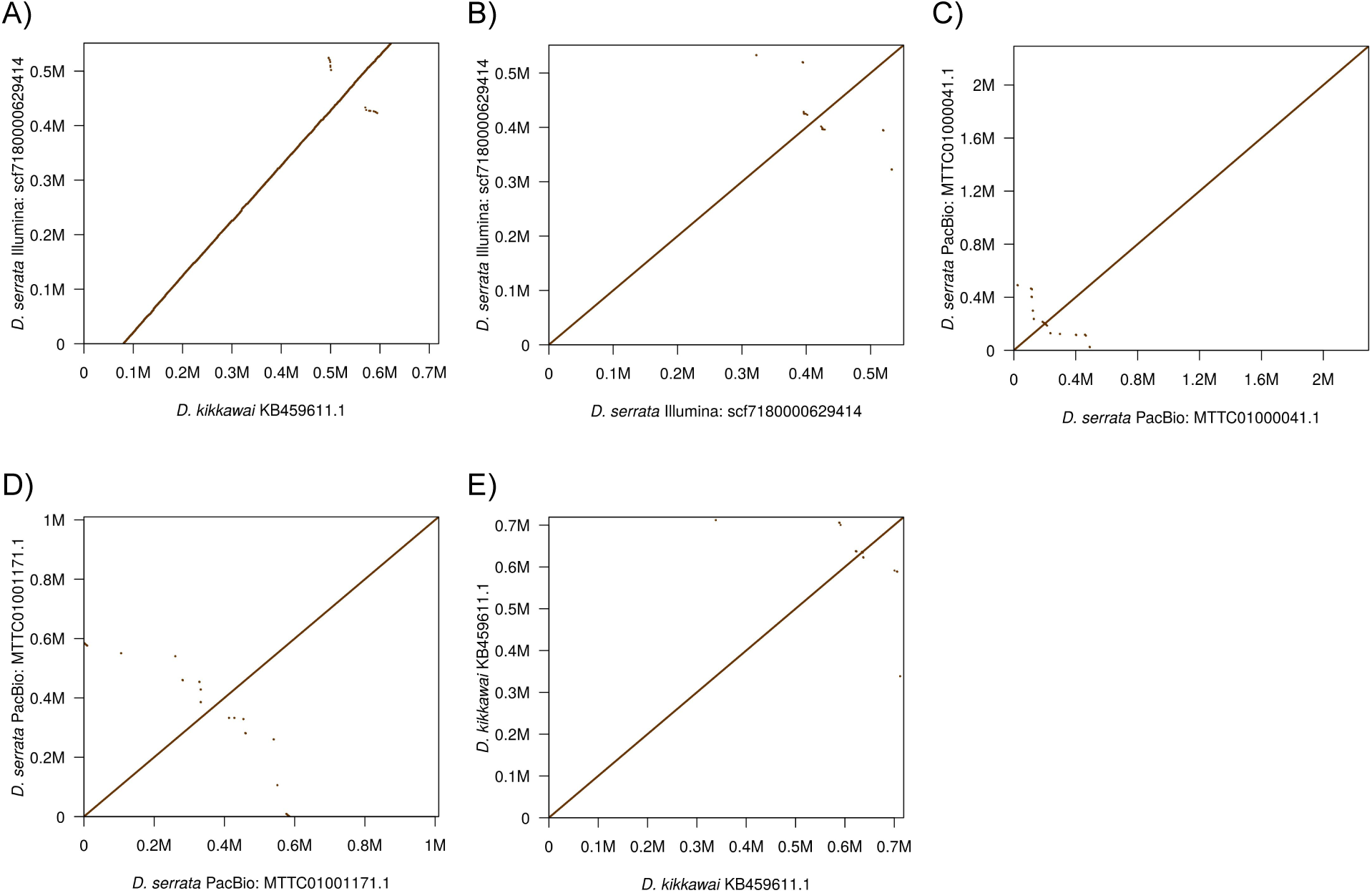
Additional dotplots. A) The alignment of the fifth longest scaffold (scf7180000629414) from our Illumina *D. serrata* assembly (strain 14028-0681.02) to the orthologous scaffold from the previously published *D. kikkawai* assembly [3]. The alignment is highly collinear, and our scaffold aligns end-to-end within the longer *D. kikkawai* scaffold. B) The alignment of scf7180000629414 to itself. C) and D) The alignment of contigs MTTC01000041.1 and MTTC01001171.1 from the previously published *D. serrata* assembly (strain Fors4) [19] to themselves. Portions of these contigs aligned to scf7180000629414. E) The alignment of scaffold KB459611.1 from the *D. kikkawai* assembly [3] to itself. This is the same *D. kikkawai* scaffold from Part A). All pairwise alignments were generated by LASTZ [39].

